# Bacillus Calmette-Guerin infection in NADPH oxidase deficiency: defective mycobacterial sequestration and granuloma formation

**DOI:** 10.1101/005835

**Authors:** Christine Deffert, Michela G. Schäppi, Jean-Claude Pache, Julien Cachat, Dominique Vesin, Ruth Bisig, Xiaojuan Ma Mulone, Tiina Kelkka, Rikard Holmdahl, Irene Garcia, Maria L. Olleros, Karl-Heinz Krause

## Abstract

Patients with chronic granulomatous disease (CGD) lack generation of reactive oxygen species (ROS) through the phagocyte NADPH oxidase NOX2. CGD is an immune deficiency that leads to frequent infections with certain pathogens; this is well documented for *S. aureus* and *A. fumigatus*, but less clear for mycobacteria. We therefore performed an extensive literature search which yielded 297 cases of CGD patients with mycobacterial infections; *M.bovis* BCG was most commonly recovered (74%). The relationship between NOX2 deficiency and BCG infection however has never been studied in a mouse model. We therefore investigated BCG infection in three different mouse models of CGD: Ncf1 mutants in two different genetic backgrounds and NOX2 knock-out mice. In addition we investigated a macrophage-specific rescue (transgenic expression of Ncf1 under the control of the CD68 promoter). Wild type mice did not develop severe disease upon BCG injection. In contrast, all three types of CGD mice were highly susceptible to BCG, as witnessed by a severe weight loss, development of hemorrhagic pneumonia, and a high mortality (∼ 50%). Rescue of NOX2 activity in macrophages restored BCG resistance, similar as seen in wild-type mice. Granulomas from mycobacteria-infected wild type mice generated ROS, while granulomas from CGD mice did not. Bacterial load in CGD mice was only moderately increased, suggesting that it was not crucial for the observed phenotype. CGD mice responded with massively enhanced cytokine release (TNF-α, IFN-γ, IL-17 and IL-12) to BCG infection, which might account for severity of the disease. Finally, in wild-type mice, macrophages formed clusters and restricted mycobacteria to granulomas, while macrophages and mycobacteria were diffusely distributed in lung tissue from CGD mice. Our results demonstrate that lack of the NADPH oxidase leads to a markedly increased severity of BCG infection through mechanisms including increased cytokine production and impaired granuloma formation.

## Author summary

The vaccine *Mycobacterium bovis* BCG is administrated to prevent early age tuberculosis in endemic areas. BCG is a live vaccine with a low incidence of complications. However, local or disseminated BCG infection may occur, in particular in immunodeficient individuals. Chronic granulomatous disease (CGD), a deficiency in the superoxide-producing phagocytes NADPH oxidase, is one of the most frequent genetic immune defects in humans. Here we analyze the role of the phagocyte NADPH oxidase NOX2 in the defense against BCG. An extensive literature review suggested that BCG infection is by far the most common mycobacterial disease in CGD patients (220 published cases). We therefore studied BCG infection in several CGD mouse models showing that these were highly susceptible to BCG infection with a mortality rate of ∼50%. As compared to the wild type, CGD mice showed a markedly increased release of cytokines, an altered granuloma structure, and were unable to restrain mycobacteria within granulomas. Rescue of the phagocyte NADPH oxidase in macrophages was sufficient to protect mice from BCG infection and to sequester the mycobacteria within granulomas. Thus, superoxide generation by macrophages plays an important role for the defense against BCG infection and prevents overshooting release of proinflammatory cytokines.

## Introduction

*M. bovis* BCG (Bacillus Calmette Guérin) is an attenuated strain of *M. bovis*, used as a vaccine against tuberculosis. BCG vaccination has a proven efficacy only early in life (<1 year of age), in particular against tuberculous meningitis and miliary tuberculosis. Thus, the WHO recommends vaccination of newborns in endemic areas. However, BCG is a live vaccine, which may persist and become a pathogen. In some individuals, in particular those with immune defects, BCG vaccination may lead to severe local or to disseminated infection [1,2]. BCG is also used as local treatment for bladder cancer [3], where in some cases it may lead to symptomatic infection, from cystitis to life threatening dissemination [4]. However, there is emerging evidence for increased risk of BCG infection in patients lacking the phagocyte NADPH oxidase (chronic granulomatous disease, CGD) [5–7]. Indeed studies looking at underlying risk factors in patients presenting with BCG infection suggest that approximately 20% of such patients suffer from CGD [8] and in many instances, BCG infection is the first manifestation of CGD [9].

The phagocyte NADPH oxidase NOX2 is a superoxide producing enzyme, involved in the host defense against numerous bacteria and fungi. Genetic loss of function of NOX2 is a primary immunodeficiency referred as chronic granulomatous disease (CGD). CGD may be caused by mutations in NOX2 or one of its subunits, in particular *p*47^phox^, which is coded by the *Ncf1* gene [10]. CGD patients suffer from severe and recurrent bacterial and fungal infections as well as from hyperinflammatory and autoimmune diseases [11]. Until about 10 years ago, it was thought that the phagocyte NADPH oxidase was not relevant for the defense against mycobacteria [12]. Whether mice carrying CGD mutations show an increase susceptibility to infection with mycobacterium tuberculosis remains controversial [13], while their susceptibility to BCG infection has so far not been studied.

Host defense mechanisms against mycobacteria are typically initiated by phagocytosis through macrophages, inducing inflammation and subsequently cell-mediated immunity involving Th1-type immune responses. These coordinated mechanisms result in granuloma formation. Granulomas are highly organized structures generated by interactions between myeloid and lymphoid cells that characterize the adaptive immune response to mycobacteria. In general granulomas sequester mycobacteria and thereby limit their dissemination. Granulomas are formed through cellular recruitment and are associated with production of cytokines and chemokines [14]. Among these cytokines, TNF and IFN-γ are the main players contributing to activation of macrophage host defense mechanism [15]. Neutrophils are able to kill mycobacteria *in vitro*, but the *in vivo* relevance of neutrophils in the mycobacterial host defense remains a matter of debate [16].

Here we have first analyzed the relevance of BCG infection in CGD patients and then investigated the role of NADPH oxidase-generated ROS in experimental BCG infection. Mice lacking a functional phagocyte NADPH oxidase showed a markedly enhanced severity to BCG infection. Rescue of phagocyte NADPH oxidase function in macrophages was sufficient to reverse the phenotype to the mild disease observed in wild-type mice. We identified increased cytokine generation and poorly organized granuloma formation as mechanisms involved in the exacerbated severity of BCG infection in NADPH oxidase-deficient mice.

## Results

To understand the relevance of mycobacterial infections for CGD patients, we performed an extensive review of the existing studies and case reports on this topic. A previous literature-based study from 1971 to 2006 reported 72 cases [31]; here we report a total of 297 cases of mycobacterial infections in CGD patients [8,31–38]. *M. bovis* BCG infection was by most frequently reported (74%; i.e. 220 cases); 20% of cases were caused by *Mycobacterium tuberculosis* infection, and the rest by different nontuberculous mycobacteria (Fig. 1A). Most of the BCG cases were local or regional infections. Relatively little information was found on the treatment and the outcome of these local infections, however one article mentions the necessity for surgical excision [31]. However, systemic BCG infections in CGD patients were not uncommon (31 cases; Fig. 1B). In 6 of the 31 cases, the outcome has been documented: 3 of the patients died and 3 of the patients survived [31,39–43].

**Figure 1.**
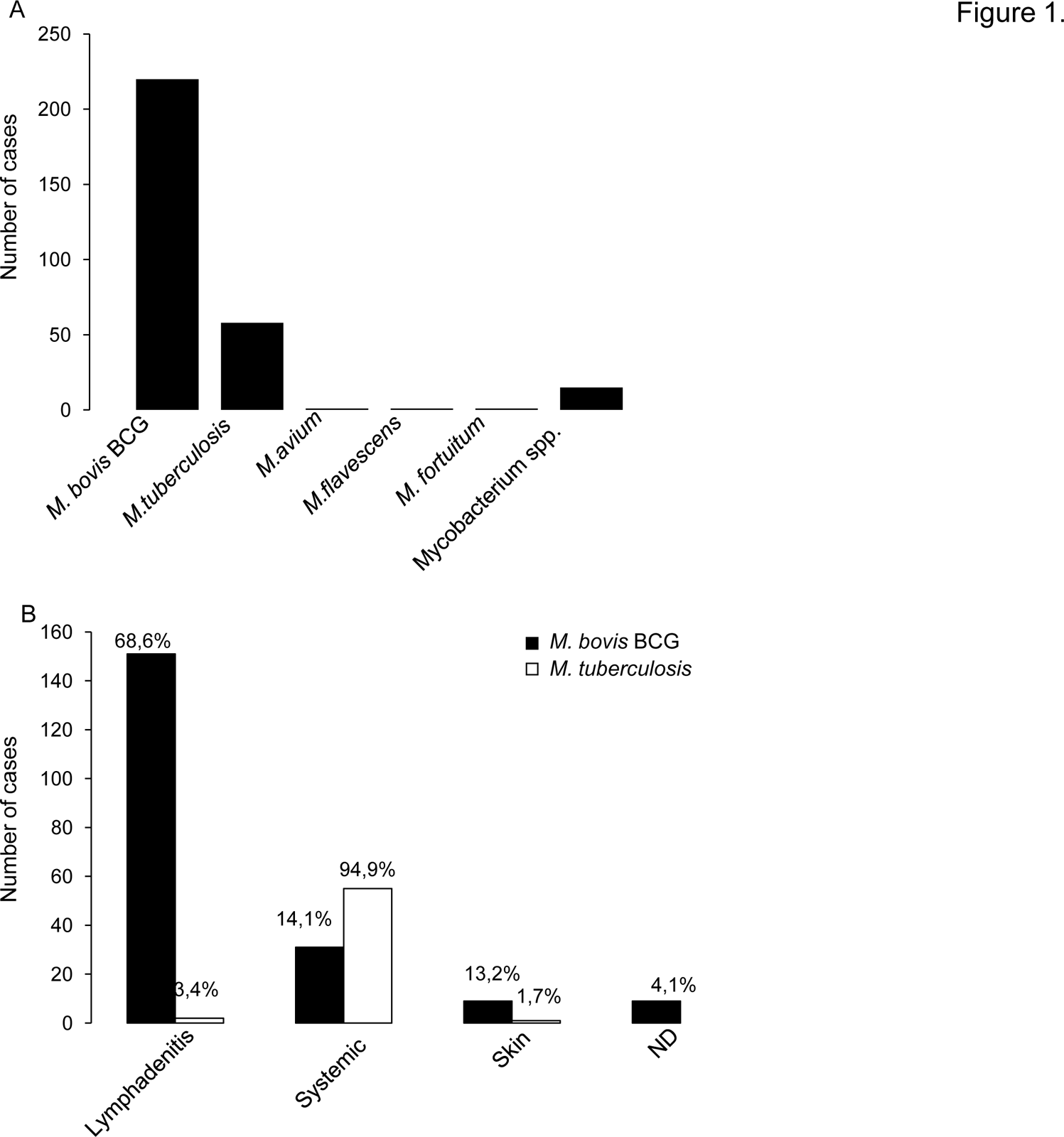
Analysis of published cases of mycobacterial infections in CGD patients. Our literature research identified a total of 297 published cases of mycobacterial disease in CGD patients. (A) Mycobacterial species recovered in mycobacterial disease in CGD patients. (B) Clinical presentations of *Mycobacterium bovis* BCG and *Mycobacterium tuberculosis* infections in CGD patients. The numbers indicated on top of each column represent the percentage with respect to the total number of BCG or M. tuberculosis cases, respectively. The terms “systemic” refers to disseminated or to lung infections.

We therefore studied BCG infection in CGD mouse models. To exclude epistatic effects, we investigated different types of CGD mice (Ncf1 mutants, NOX2-knock-out mice), as well as different genetic backgrounds (C57Bl/10.Q, C57Bl/6). Taken together, the following mouse lines were used:

i. wild type C57Bl/10.Q mice;
ii. Ncf1 mutant mice with a loss of function mutation in the p47phox subunit of the phagocyte NADPH oxidase (genetic background: C57Bl/10.Q; [22,44]).
iii. Ncf1 mutant mice with a transgenic rescue of the Ncf1 gene under the control of a CD68 promoter. These mice express a wild-type Ncf1/p47^phox^ protein in macrophages and dendritic cells, but not in neutrophils and are referred to as *Ncf1* rescue mice (genetic background: C57Bl/10.Q) [18].
iv. Ncf1 mutant mice with C57Bl/6 background
v. NOX2-deficient mice with C57Bl/6 background [45].

## ROS production in mononuclear phagocytes limits severity of BCG infection.

Mice were injected intravenously with BCG (10^7^ CFU). Wild-type as well as *Ncf1* rescue mice resisted to the infection during the 4 weeks observation. In contrast, *Ncf1* mutant mice showed early mortality: 50% of mice died after 10 days and only 33% of the mice survived after 4 weeks (Figure 2A). The high mortality of *Ncf1* mutant mice was associated to a rapid weight loss, which was absent in wild-type controls and *Ncf1* rescue mice (Figure 2B). To examine if there was a late mortality in *Ncf1* rescue mice, survival was monitored for up to 20 weeks. Only one on three *Ncf1* rescue mice died at 11 weeks after BCG infection. To investigate whether the genetic background or the type of CGD mutation was responsible for the high susceptibility, we investigate other types of CGD mice. BCG-infected *Ncf1* mutant mice in a C57Bl/6 background showed a mortality comparable to the one observed in a.l/10Q background (Figure 2C) and similarly showed a rapid weight loss (Figure 2D). Also, BCG infection in NOX2-deficient mice led to a high mortality and weight loss as compared to wild type controls (Figure 2E and F). These observations strongly suggest that in absence of NADPH oxidase, mice are more susceptible to mycobacterial infection independently of the mouse genetic background. Furthermore, the almost normal survival of *Ncf1* rescue mice implies that ROS production in mononuclear phagocytes is crucial.

**Figure 2.**
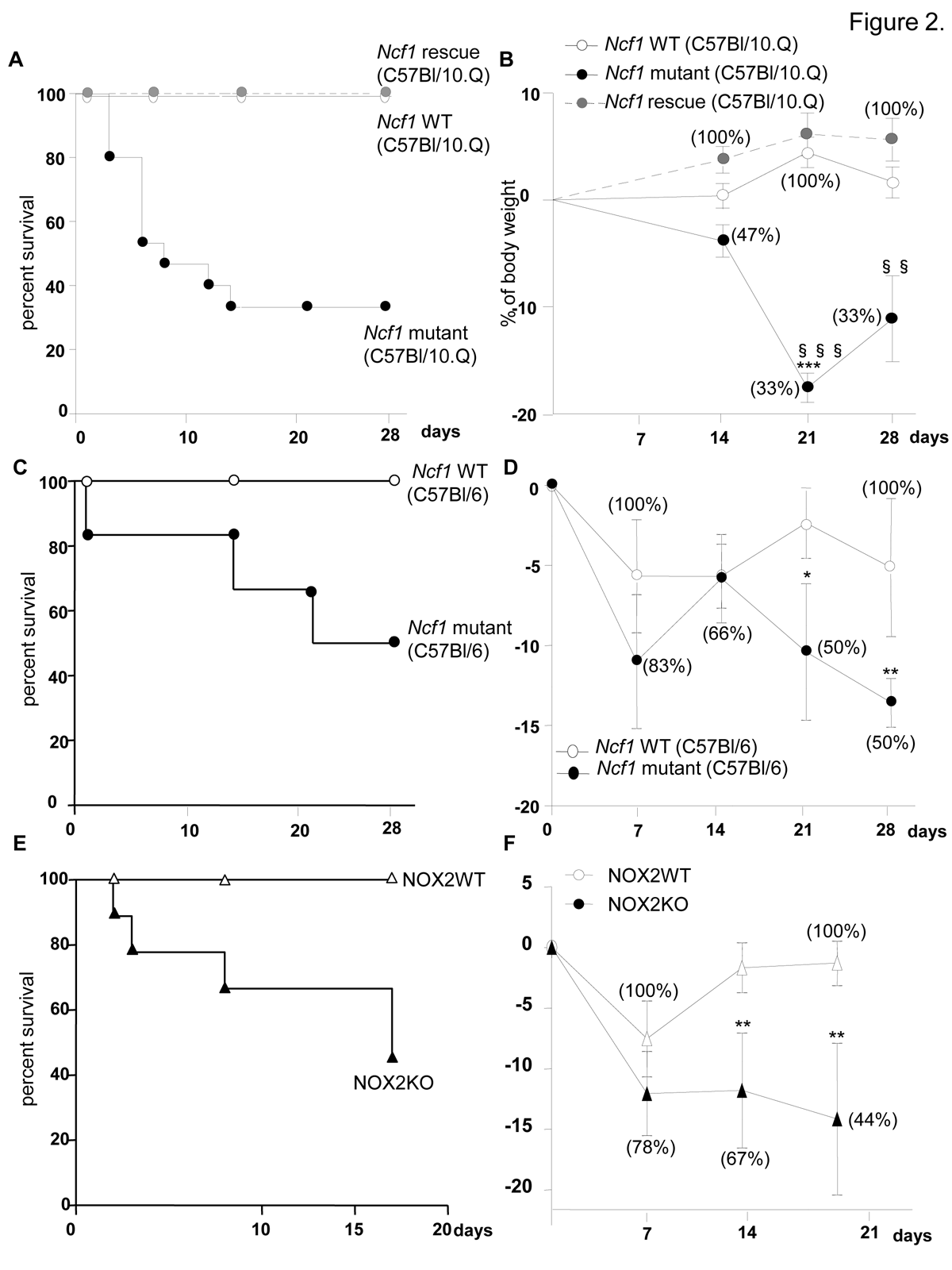
Impact of CGD mutation on mortality and weight loss in response to BCG infection. *Ncf1* mutant (loss of function mutation in p47^phox^), *Ncf1* rescue (expression of wild-type p47^phox^ in mononuclear phagocytes) with C57Bl/10.Q background, *Ncf1* mutant (loss of function mutation in p47^phox^) with C57Bl/6 background, NOX2-deficient and their respective wild-type controls were injected intravenously with BCG (10^7^ CFU). Survival was monitored over the 4 weeks period following BCG inoculation in (A) C57Bl/10.Q wild-type (n = 15), *Ncf1* mutant (n = 15) and *Ncf1* rescue (n = 11), in (C) C57Bl/6 wild-type (n = 7), *Ncf1* mutant (n = 6) and (E) C57Bl/6 wild-type (n = 7), NOX2-deficient (n = 9) mice. (B, D and F) Body weight changes as a function of time after BCG inoculation. Survival (percent of initial number of mice) is shown in brackets. Statistics shown in the figures are the comparison between respective wild-type and *Ncf1* mutant or NOX2-deficient (***: p < 0.001, **: p < 0.05, *: p < 0.01) and the comparison between *Ncf1* mutant and rescue (§§§: p < 0.001 and §§: p < 0.01). Note that no significant differences were observed between wild-type and *Ncf1* rescue mice.

## Severe pulmonary lesions after BCG inoculation in the phagocyte NADPH oxidase-deficient mice.

To further understand the causes of the early mortality of *Ncf1* mutant mice, mice were sacrificed at day 3 post-infection for lung histopathological examination. In the absence of BCG infection, no histological differences were observed between wild-type, *Ncf1* mutant and rescue mice (Supplementary Figure S1). However, upon BCG infection, *Ncf1* mutant mice presented severe inflammatory lesions in the lungs with extended hemorrhagic lesions, intravascular thrombosis, decrease of the alveolar spaces (Figure 3A.b) and severe pleurisy (Figure 3B.b). *Ncf1* mutant mice also showed accumulation of inflammatory cells composed essentially by neutrophils concentrated as microabscesses (Figure 3A.e). In contrast, wild-type and *Ncf1* rescue did not show massive hemorrhagic lesions and only moderate inflammation with mixed inflammatory cells observed in lungs (Figures 3A.a, c, d and f) and pleura (Figures 3B.a and c).

**Figure 3.**
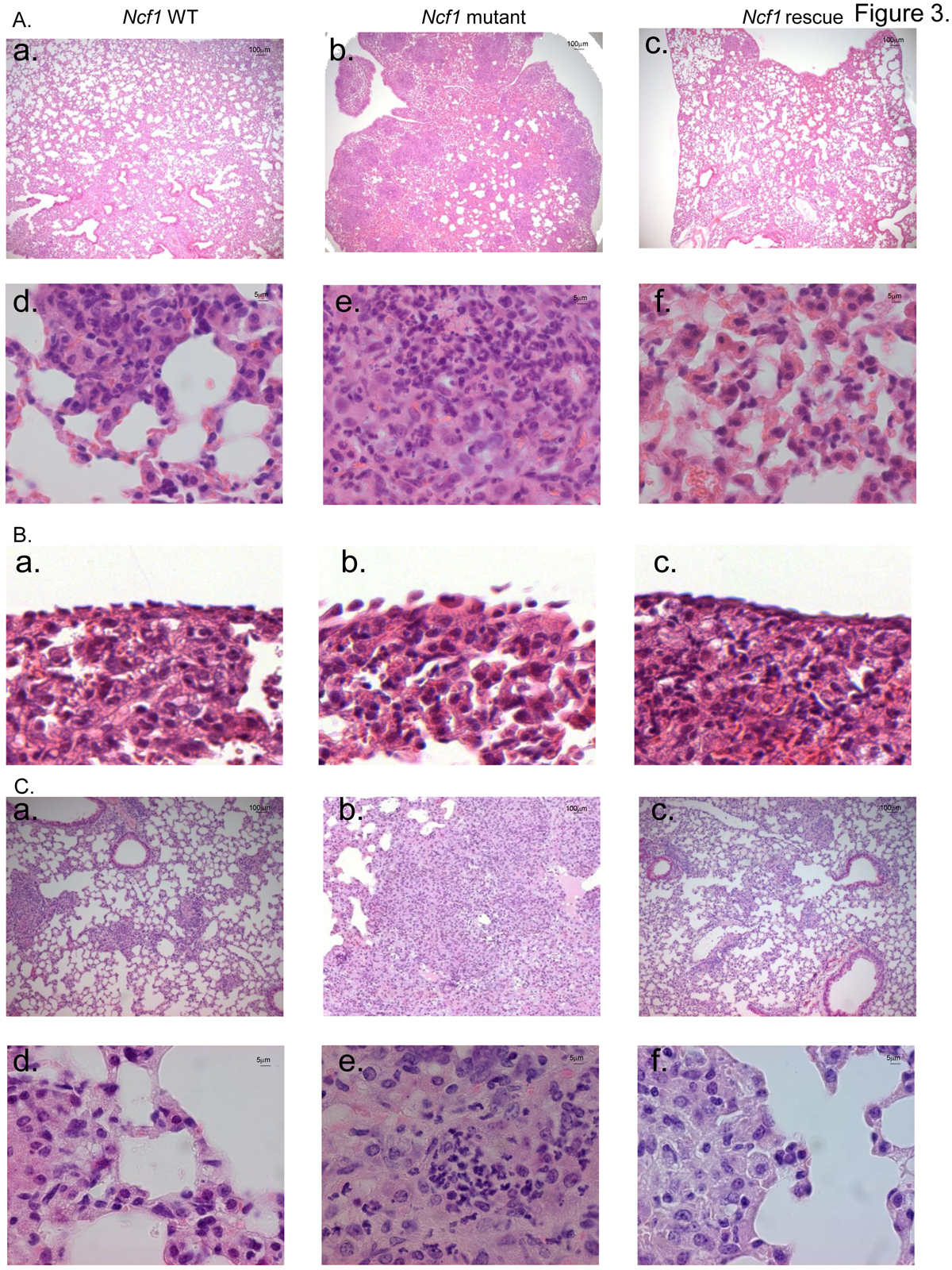
*Ncf1* mutation leads to severe lung damage in response to BCG infection. Hematoxylin and eosin-stained lung sections taken from mice sacrificed after 3 days (A, B) or 4 weeks (C) of BCG infection. (A, C) representative sections from wild-type (a and d), *Ncf1* mutant (b and e) and *Ncf1* rescue (c and f) mice. (B) Pleural histology from wild-type (a), *Ncf1* mutant (b) and *Ncf1* rescue (c) mice sacrificed 3 days after BCG infection. Magnifications were x200 (panels A a-c and C a-c) and x1000 (panels A d-f and C d-f) and x600 (panel B).

The surviving *Ncf1* mutant mice (5 out of 15) were also analyzed at 4 weeks post-infection. Histopathological examination of *Ncf1* mutant lungs revealed extensive inflammatory lesions reducing notably the alveolar space (Figure 3C.b) and microabscesses composed of neutrophils. Only one third of the *Ncf1* mutant mice survived up to 4 weeks and the latter results might represent a survivor effect, and not necessarily be representative for all *Ncf1* mutant mice. In contrast, wild-type and *Ncf1* rescue did not show massive infiltrate of inflammatory cells (Figures 3C.a-c and d–f).

In the absence of BCG infection, the organ weight indexes for lung, liver and spleen were comparable in all mouse strains (Supplementary Figure S1). After BCG infection, lung weight, as a surrogate measure of lung inflammation and edema, increased only moderately in wild-type and *Ncf1* rescue mice, but massively in *Ncf1* mutant mice (Figure 4A). The severity of lung pathology was also assessed by analysis of free alveolar space vs. occupied space. The occupied space was significantly increased in *Ncf1* mutant as compared to wild-type and *Ncf1* rescue lungs (Figure 4B). This quantification corroborates with the massive obstruction of alveolar space in *Ncf1* mutant mice seen in histology. Similar as seen for *Ncf1* mutant mice, BCG-infected NOX2-deficient mice showed severe inflammatory lesions with extended hemorrhagic lesions and decreased alveolar space (Figure S2.A and B.b). Microabscesses of neutrophils were also present in NOX2-deficient and Ncf1 mutant in C57Bl/6 background lungs while mixed inflammatory cells were observed in wild-type lungs (Figure S2.B.d and C). The lung weight was also increased in NOX2-deficient and Ncf1 mutant in C57Bl/6 background mice compared to wild-type (Figure S3.A). In summary, based on histopathological examination, weight index, and quantification of occupied/free space in the lung, our data support a protective role of ROS production by mononuclear phagocytes in BCG infection.

**Figure 4.**
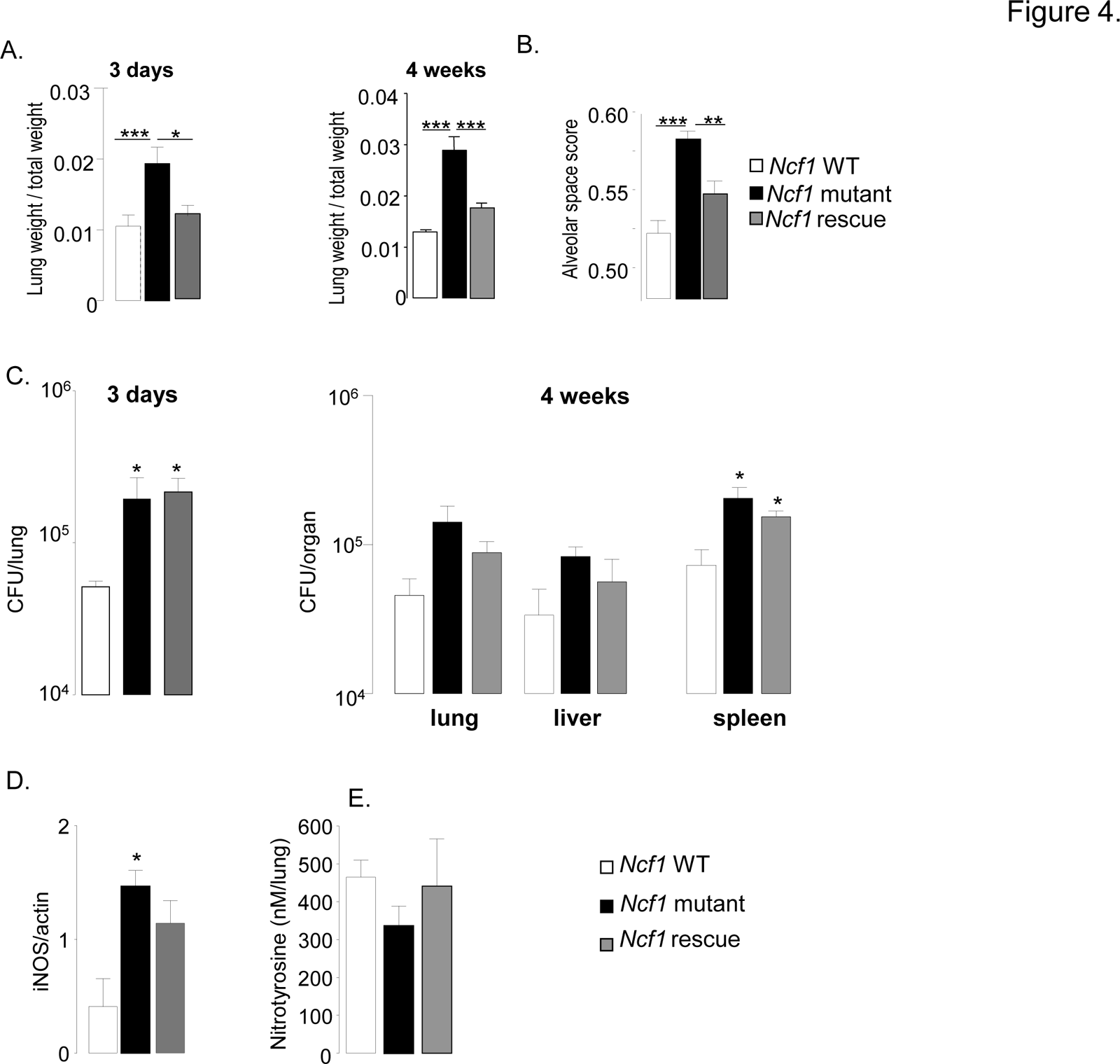
Lung parameters in response to BCG infection. (A) Lung/weight ratio of wild-type (n = 9), *Ncf1* mutant (n = 5) and *Ncf1* rescue (n = 8) mice at 3 days and 4 weeks post infection. (B) Determination of alveolar space score (occupied lung tissue *vs.* free space) in lung sections. Data are represented as the mean of alveolar space score ± SD in 4 mice per group with at least 3 lobes analyzed per mouse. (C) Number of viable bacteria was determined at 3 days and 4 weeks following BCG infection. Data are shown as mean log of CFU per organ (±SEM; 3-5 mice per group). (D) iNOS protein expression in lung was detected by western blot 4 weeks after BCG infection. Results are expressed as mean ± SEM of relative units of iNOS/actin (n = 4, per group) after quantification by Image Quant software. Nitrotyrosine quantification by ELISA was done in lungs, 4 weeks after BCG infection. Results are expressed as mean ± SEM of nM per lung (n = 4-5, per group). (***: p < 0.001, **: p < 0.05, *: p < 0.01).

## Bacterial counts and iNOS activation upon BCG infection

We quantified bacterial burden in different organs. Bacterial load in the lung of both the *Ncf1* mutant and *Ncf1* rescue mice were similar but higher than those in wild-type mice 3 days after infection (Figure 4C). Four weeks post-infection, no significant differences in bacterial load were observed in lung and liver. However, bacterial counts in the spleen of *Ncf1*mutant and *Ncf1*rescue mice were significantly increased (Figure 4C). Thus, NOX2 activity of neutrophils (not reconstituted in the rescue mice) might be involved in the early clearance of mycobacteria; note however that this effect was rather modest (∼0.6 log). We further evaluated inducible nitric oxide synthase iNOS, which is crucial for clearance of BCG and mouse survival [46]. Expression of iNOS protein in the lung at 4 weeks post-infection was significantly increased in *Ncf1* mutant mice (Figure 4D). NO and superoxide form peroxynitrite, a highly reactive molecule, are implicated in mycobacteria killing. Upon interaction with proteins, peroxynitrite produces nitrotyrosine, which are stable biological peroxynitrite markers [47]. Interestingly, despite the increase of iNOS protein, nitrotyrosine levels in lungs of *Ncf1* mutant were not different from wild-type (Figure 4D), presumably because *Ncf1* mutant mice lack the second substrate required for peroxynitrite generation, namely superoxide. Thus, most likely CGD mice produce increased amounts of NO in response to mycobacteria, but given the lack of NOX2-generated superoxide, this is not accompanied by an increase in peroxynitrite.

## Cytokine and chemokine levels activated by BCG infection

We next measured levels of selected cytokines in lung homogenates from BCG infected mice (Figures 5). Three days post infection, the increase of TNF levels in *Ncf1* mutant lung was massive (3.6-fold compared to wild-type) and also observed at four weeks post-infection (Figure 5A). TNF levels in *Ncf1* rescue lung were comparable to those observed in wild-type lung. Three days, but not 4 weeks, post infection, IL-17 lung levels were increased in *Ncf1* mutant mice (Figure 5B). The same pattern was observed for IL-12p40 (Figure 5C). The pattern was slightly different for IFN-γ: there were increased in *Ncf1* mutant mice at 3 days and 4 weeks post-infection, however *Ncf1* rescue mice showed even higher IFN-γ levels 4 weeks post-infection (Figure 4D). We also assessed the levels of the chemokines CXCL1 (KC, the murine IL-8 homolog), and CCL5 (RANTES). We selected CXCL1 because it is a powerful neutrophil chemoattractant and might explain the high number of neutrophils in the lung lesion in mutant mice and CCL5 because it is a leukocyte chemoattractant with a potentially important role in granuloma formation. *Ncf1* mutant mice showed a higher CXCL1 levels 3 days after BCG infection (Figure 5E). CCL5 levels were increased in *Ncf1* mutant mice three days and 4 weeks after infection (Figure 5F). The general pattern was an increase in pulmonary cytokine and chemokine responses in *Ncf1* mutant mice due to the infection which was controlled by *Ncf1* rescue mice. Moreover, we also evaluated if NOX2-deficient mice would also respond with an exacerbated cytokine response using ex-vivo recall of spleen cells from BCG infected mice. Both re-infection of splenocytes or addition of BCG antigens resulted in enhanced TNF and nitrite, as an indicator of NO production confirming that NOX2 deficiency leads to an increase response in TNF and immune mediators (Supplementary Figure S3).

**Figure 5.**
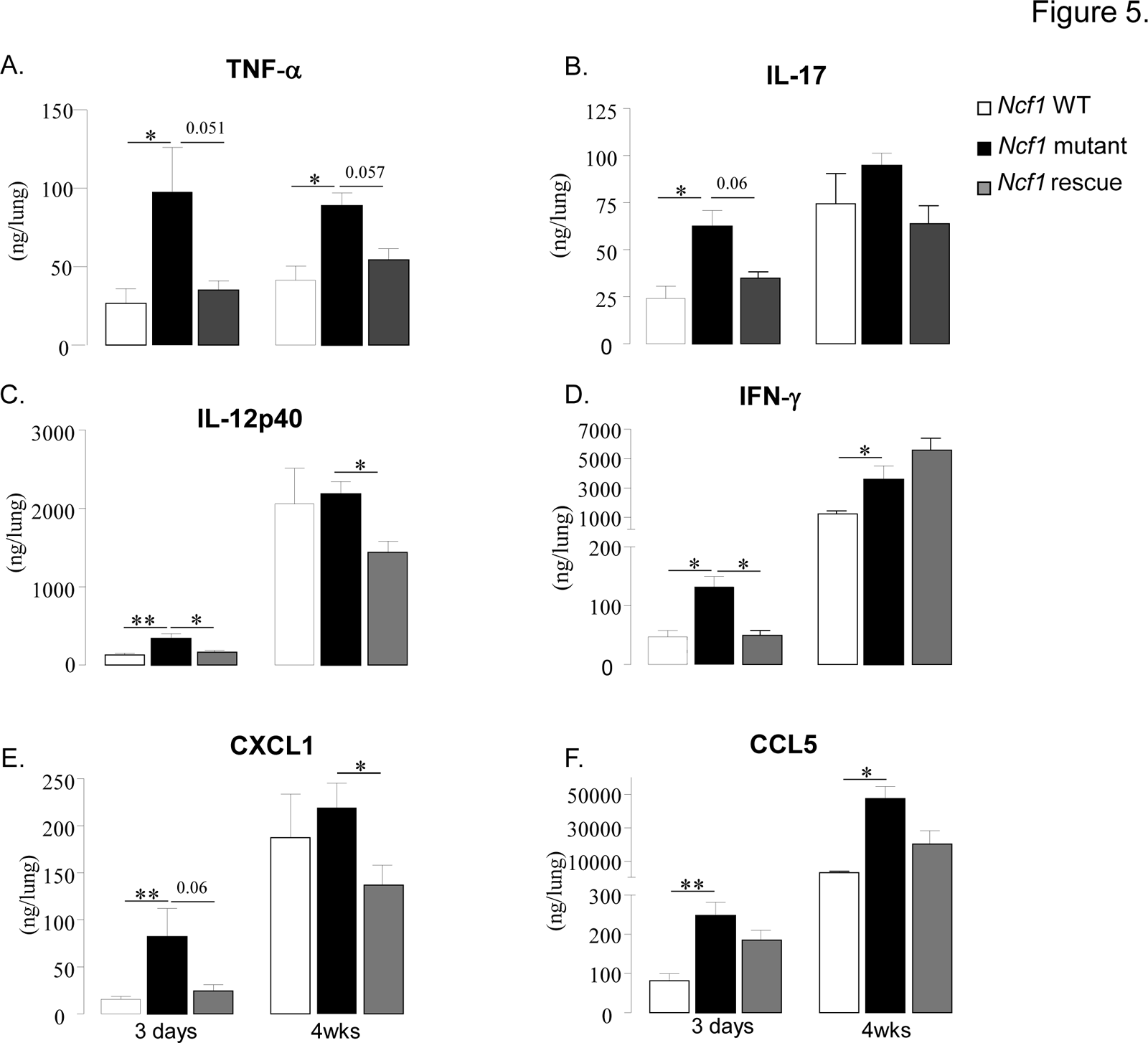
Cytokine and chemokine responses to BCG infection. TNF-α (A), IL-17 (B), IL-12p40 (C), IFN-γ (D), CXCL1 (E) and CCL5 (F) were assessed in lung homogenates obtained 3 days and 4 weeks after BCG infection. Results are presented as the mean ± SEM (n = 4-7 mice per group). (**: p < 0.05, *: p < 0.01).

## ROS generation in mononuclear cells and in granulomas from *Ncf1* rescue mice

We have previously demonstrated that ROS production in response to phorbol myristate acetate (PMA) or β-glucan was abolished in neutrophils, macrophages and dendritic cells in *Ncf1* mutant mice [22]. We therefore investigated ROS production in macrophages and dendritic cells exposed to BCG. Wild-type and *Ncf1* rescue cells produced ROS in response to BCG, but not *Ncf1* mutant cells (Figures 6A, B). Diphenylene iodonium (DPI), a non-specific NOX inhibitor, abolished the mycobacteria-induced ROS production in wild-type and *Ncf1* rescue cells. The kinetic of ROS production was comparable in macrophages and dendritic cells from wild-type and *Ncf1* rescue mice (Figures 6A and B).

**Figure 6.**
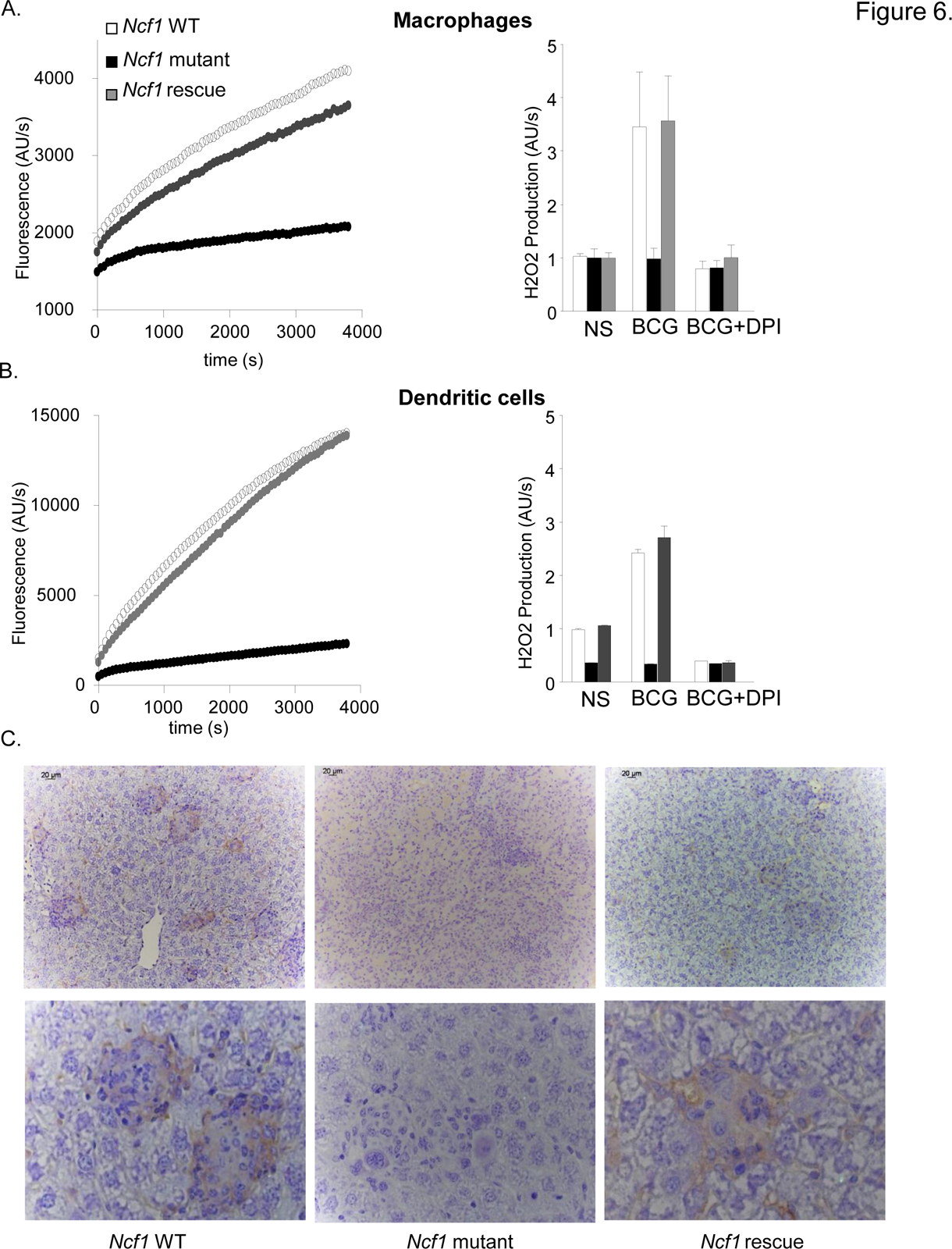
ROS generation in response to BCG *in vitro* and *in vivo.* (A) Bone marrow-derived macrophages and (B) dendritic cells were obtained from wild-type, *Ncf1* mutant and *Ncf1* rescue mice. H2O2 release was measured by Amplex red. Left panels: Cells were stimulated with BCG. Representative kinetic graphs of ROS production as measured by fluorescence emission. Right panels: Cells were exposed to PBS (NS) or BCG; where indicated, the inhibitor DPI was added. Histograms representing ROS production of three independent experiments performed in duplicate (mean ± SEM). (C) Representative images of immunohistochemistry for 8-hydroxy-2′-deoxyguanosine (8-OHdG) in liver sections from wild-type, *Ncf1* mutant and rescue mice obtained 4 weeks post infection. Magnifications were x200 (upper panel C) and x1000 (lower panel).

We next investigated whether there are signs of ROS production in granulomas *in vivo*. For this purpose, liver sections from 4 weeks BCG-infected mice were stained with an antibody against 8-OHdG (8-hydroxydeoxyguanosine), a well-studied marker of DNA oxidation. In wild-type and *Ncf1* rescue liver, an important 8-OHdG staining was observed within granulomas (Figure 6C). Note that, to the best of our knowledge, this is the first demonstration of ROS generation during granuloma formation. Importantly, no 8-OHdG staining was observed in granulomas of *Ncf1* mutant mice, demonstrating that the phagocyte NADPH oxidase is the major source of ROS during BCG infection.

## Phagocyte NADPH oxidase in mononuclear phagocytes contributes to granuloma formation and sequestration of mycobacteria

We next analyzed lung histology 3 days and 4 weeks after BCG infection using the following stainings: H/E (general morphology), Ziehl-Neelsen (mycobacteria), acidic phosphatase activity (activated macrophages). Three days post BCG infection, lung sections from wild-type and *Ncf1* rescue mice showed clusters of macrophages (Figure 7A). The reaction appeared different in *Ncf1* mutant mice: BCG infection induced abundant neutrophil abscesses, with a lack of macrophage clustering within restricted areas (Figure 7B). Mycobacteria appeared less abundant in lung section from wild-type as compared to *Ncf1* mutant mice (Figure 7C). Interestingly, the Ziehl-Neelsen stain suggests a relatively high bacterial load in rescue mice, corroborating the quantitative bacterial load analysis (Figure 4C).

**Figure 7.**
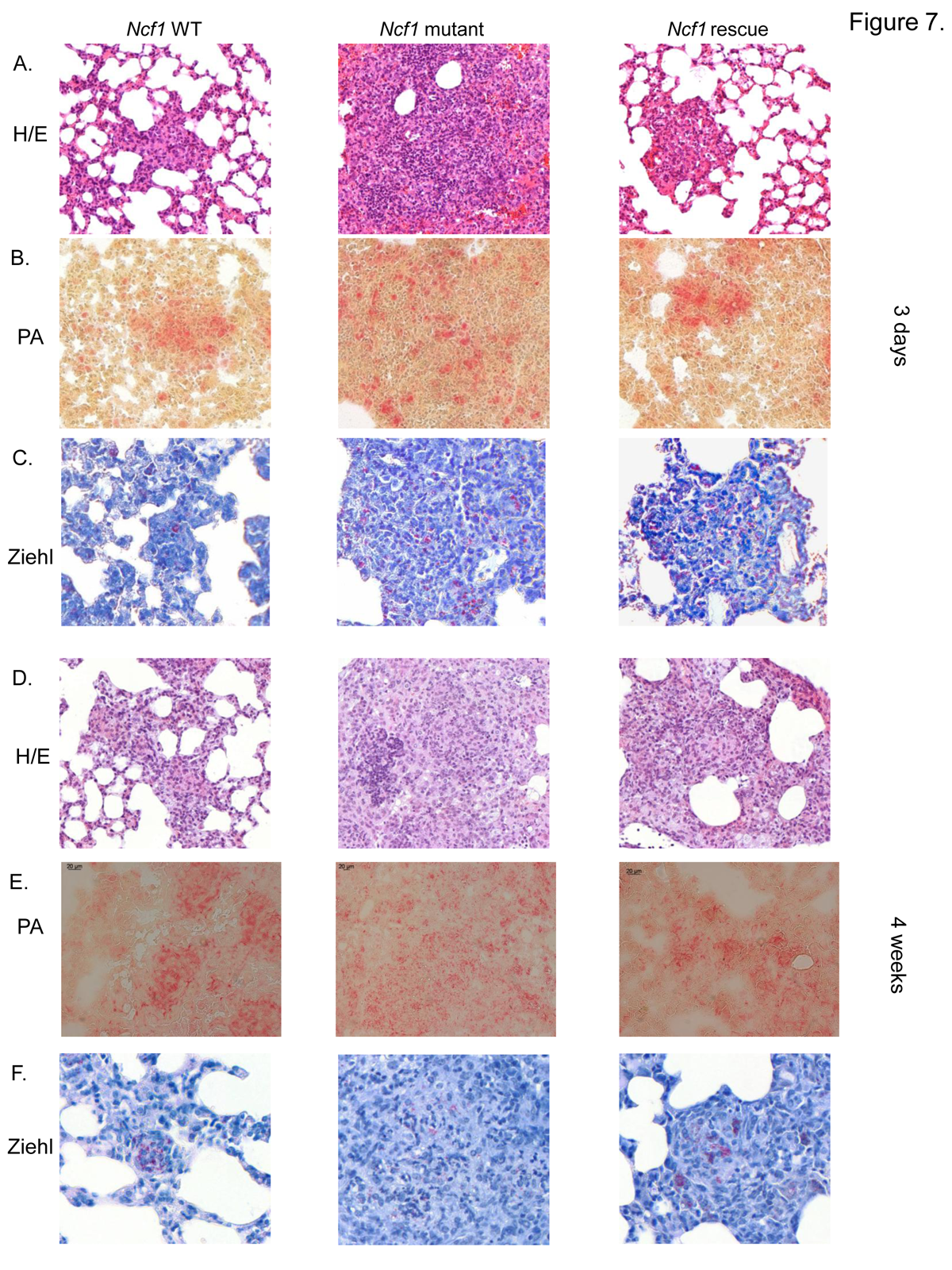
Role of the phagocyte NADPH oxidase in granuloma formation and BCG sequestration. Three days and 4 weeks after BCG infection, wild-type, *Ncf1* mutant and rescue mice were sacrificed and lungs were fixed and sectioned. Representative hematoxylin and eosin (H&E) stained lung histology focused on (A) focal clusters at 3 days and (D) granuloma formation at 4 weeks. Representative acid phosphatase activity in lung lesions showing activated macrophages at 3 days (B) and 4 weeks (E). Ziehl-Neelsen staining of lung lesions showing distribution of acid fast bacilli at 3 days (C) and 4 weeks (F). Magnifications were x200.

Granuloma formation is a crucial step in the mycobacterial containment and clearance. After 4 weeks of BCG infection, both wild-type and *Ncf1* rescue mice had well differentiated granulomas containing multinucleated giant cells (Figures 7D–E). In these mice, granulomas enclosed the mycobacteria and virtually no mycobacteria were observed outside of granulomas (Figure 7F). In contrast, *Ncf1* mutant mice presented large pyogranulomatous lesions with abundant neutrophil abscesses (Figure 7D) and diffusely distributed acid phosphatase-positive macrophages (Figure 7E). Importantly, no sequestration of mycobacteria was observed (Figure 7F). As seen for *Ncf1* mutant mice, NOX2-deficient mice as well as *Ncf1* mutant mice in a C57Bl/6 background showed larger granulomas without concise delimitations (Supplementary Figure S4B). Thus, the presence of NADPH oxidase in mononuclear phagocytes is required for the formation of compact granulomas with concise delimitations and for sequestration of mycobacteria within granulomas.

## Discussion

In this study, we have analyzed BCG infection in several mouse model of phagocyte NADPH oxidase deficiency. CGD mice were highly susceptible to BCG infection. Our results suggest that the phagocyte NADPH oxidase limits the severity of mycobacterial infection by at least two mechanisms: i) block of overshooting cytokine release; and ii) contribution to mycobacterial sequestration in granulomas. For these two mechanisms, NADPH oxidase function in macrophages was essential. We also observed a modest increase in bacterial load in mice lacking NOX2 function, an effect which appeared to depend on NOX2 in neutrophils. <u>Functionally relevant NOX2 expression in mycobacterial infection: neutrophils vs. macrophages.</u> The *Ncf1* mutant mice used in this study, including a selective rescue in mononuclear phagocytes, provide a unique experimental system to study a cell type-specific role of NOX2 in the defense against BCG [18]. Generally speaking, most of the enhanced mycobacterial pathology associated with NOX2-deficiency (morbidity, mortality, enhanced cytokine production, abnormal granuloma formation) can be attributed to macrophages. There is one exception to this: the increased mycobacterial load, which is not reversed by *Ncf1* rescue in macrophages and hence was not correlated to the outcome of infection. Particularly interesting in this respect is the recent discovery of a family with a peculiar variant of CGD [32]. These patients lack ROS production in macrophages, but not in neutrophils and showed a high sensitivity to mycobacterial infection, in particular to BCG. Thus, while our results in mice show that selective rescue of NOX2 in macrophages restores resistance BCG infection; in these CGD patients a selective loss of NOX2 in macrophages establishes high susceptibility.

<u>Role CGD mutation and genetic background for the increased severity of BCG infection in the absence of the phagocyte NADPH oxidase</u>. Hitherto, BCG infection has never been investigated in mouse models of CGD. However, infection of CGD mice with other types of mycobacteria led to discordant results: in some cases aggravation was observed [48–50], while in other studies no effect was observed [13,51,52]. It seems plausible that for infections with virulent mycobacteria, such as *M. tuberculosis*, the role of the phagocyte NADPH oxidase is less relevant [13]. Yet, we wanted to assure that our results are not due to a specific choice of the CGD mutation or to the genetic background. We therefore tested two different CGD mutations (*Ncf1*, *Cybb*/NOX2) as well as two different genetic backgrounds (C57/B10.Q, C57Bl/6). All results concur: CGD mice are highly susceptible to BCG infection.

<u>Enhanced neutrophil infiltration in the absence of the phagocyte NADPH oxidase</u>. Reconstitution of the phagocyte NADPH oxidase in mononuclear phagocytes completely reversed the neutrophil influx phenotype. Thus, it is not the lack of activity of NOX2 in neutrophils which leads to the increased number of neutrophils in inflammation. Most likely, NOX2 in mononuclear phagocytes regulates the number of invading neutrophils by controlling the release of neutrophil chemoattractants. These chemoattractants might be directly released from macrophages or possibly from other cells that depend on a macrophage signal. An alternative theory is the decreased uptake of apoptotic neutrophils by NADPH oxidase-deficient macrophages [53].

<u>Neutrophil NOX2 appears to limit the multiplication of BCG</u>. Our results show a small, but statistically significant increase of mycobacterial load in *Ncf1* mutants (∼0.6 log). It is unlikely that this is an important factor in the increased morbidity of BCG-infected CGD mice, as the *Ncf1* rescue mice did well, but had the same bacterial load as the Ncf1 mutants. It is however of interest that the increased mycobacterial load is not reversed by the selective rescue of NADPH oxidase in mononuclear cells. This suggests that NADPH oxidase in neutrophils has a modest, but significant contribution to killing of BCG during early infection. Indeed it has been previously suggested that neutrophils limit mycobacterial survival during early infection [16,54,55]. Note that NADPH-oxidase dependent killing of mycobacteria does not necessarily signify a direct antibacterial action of ROS. Indeed, mycobacteria have been suggested to be sensitive to neutrophil extracellular trap (NET) [56] and NADPH oxidase-dependent NET formation [57] could also be a relevant mechanism how neutrophils influence the multiplication of mycobacteria.

<u>Cytokine production in response to BCG infection</u>

It has been suggested that the increased sensitivity of CGD patients to mycobacterial infection might be linked to a ROS activation of cytokine production, in particular IL-12 (which is secreted by macrophages to stimulate IFN-γ release by T lymphocytes [2]). In CGD patients, an hyperresponsiveness of neutrophils to different stimuli was usually observed [58]. In our study, we observed the opposite: CGD mice infected with BCG generated increased levels of cytokines. Interestingly, several of the cytokines increased in *Ncf1* mutant mice (in particular TNFα, IL-12, IFN-γ, IL-17) are involved in the antimycobacterial defense. This might be a defense mechanism compensating for the lack of ROS generated by the NADPH oxidase. However, the high level of these cytokines in CGD mice might also account for the increased mortality and absence of resolution of inflammation to mycobacterial infections.

<u>Macrophage NOX2 and granuloma formation.</u>

Our results shed new light on granuloma formation in mycobacterial infection and the role of NOX2 in this process:

i. Within mycobacteria-induced granulomas there is an oxidative environment, which is completely abolished in *Ncf1* mutant mice demonstrating that NOX2 is the source of oxidative stress. Rescue of functional NOX2 in mononuclear cells was sufficient to restore the oxidative environment within the granulomas.
ii. The absence of macrophage NOX2 leads to morphologically altered granulomas. BCG-induced granulomas in *Ncf1* mutant mice were larger, but of atypical appearance: acidophilic centers were not detectable; numerous neutrophils were infiltrated; and the delimitation of granuloma boarders was barely perceivable. At early time points, macrophage clusters, presumably the earliest signs of granuloma formation, were detected in wild-type and rescue, but not in *Ncf1* mutant mice.
iii. *Ncf1* mutant granulomas are not capable of sequestering BCG. Indeed, while large, concentrated clusters of mycobacteria were readily detected in wild-type and in rescue mice, the bacteria were widely distributed throughout the tissue in *Ncf1* mutant mice. Thus, while - at least at later time points - the allover mycobacterial load was not different between *Ncf1* mutant and rescue mice, their tissue distribution was fundamentally altered.

Taken together, our results provide strong evidence for a role of NADPH oxidase-dependent ROS generation in the fine tuning of granuloma formation. Thus, redox-sensitive signaling steps are involved in the coordinated genesis of granulomas, and the overshooting cytokine and chemokine productions observed in CGD mice probably destabilizes granulomas.

To which extend do results obtained in our study apply to CGD patients? Clearly, our analysis of the published literature demonstrates that, human CGD patients are sensitive to infection with the vaccinal BCG strain [5,7]. Approximately 15% of CGD patient with BCG disease will develop a disseminated form, also referred to as BCGosis. At this point, it is not clear which are the factors precipitating such disseminated disease. Genetic modifiers, type of BCG strain, inoculum size of viable mycobacteria are among the possible culprits. Similar as observed in our mouse model of disseminated BCG infection, there was a substantial mortality associated with BCGosis in CGD patients.

Taken together, the results presented here not only shed new light on BCG infection in CGD, but also provide first evidence for a role of the macrophage NADPH oxidase in the coordination of granuloma formation. The vaccinal BCG strain is an important tool for the control of childhood tuberculosis in countries with a high incidence of the disease. In general, children are vaccinated at birth, because the major effect of BCG vaccination is prevention from tuberculous meningitis early in life. Thus, the vaccination occurs prior to first manifestations of immune deficiency. New algorithms need to be defined to assure vaccine protection of immunocompetent neonates, without putting immunodeficient neonates at risk.

## Material and methods

### Ethics statement

Animal experiments were approved by ECAE (Ethics Committee for Animal Experimentation) for the University of Geneva and the Cantonal Veterinary Office (Authorization No. 1005/3715/2). Handling and manipulation of the animals complied with European Community guidelines (FELASA).

## Mice

Wild-types B10.Q, *Ncf1* mutant and rescue mice, backcrossed into identical background were used (for details of backcross see [17,18]). *Ncf1* Rescue mice are *Ncf1* mutant animals which contain a transgenic wild-type Ncf1 gene under the control of a human CD68 promoter fragment. *Ncf1* mutant with the same mutation on a C57Bl/6N background and its respective wild-type controls were used. NOX2-deficient mice and respective controls were backcrossed on C57Bl/6 background (Jackson Laboratories). For all experiments, mice aged 8–12 weeks were kept in a quiet room at 25 ◦C with a 12 h light/dark cycle and food and water were supplied *ad libitum*.

### Experimental infection

Mice were infected intravenously with 10^7^ living CFU of *M. bovis* BCG Connaught [19]. Mortality and body weights were monitored during infection. Three days and 4 weeks post-infection, mice were sacrificed and lung, liver and spleen were weighted, fixed and frozen for subsequent analyses.

### Determination of colony forming units (CFU) from infected organs

The number of viable bacteria recovered from frozen organs was evaluated as previously described [20,21].

### Isolation and culture of primary macrophages and dendritic cells

Bone marrow primary cells were obtained from mice by flushing both the femur and the tibia as previously described [22,23]

### ROS evaluation

BMDMs and BMDCs were stimulated with BCG (MOI 10). The production of ROS by NOX2 was measured using Amplex red (Invitrogen) fluorescence, as described previously [24].

### iNOS and nitrotyrosine quantifications

Lung homogenates were prepared and western blot performed as previously described [25]. Nitrotyrosine, a stable end product of peroxynitrite oxidation, was assessed in serum by enzyme-linked immunosorbent assay (ELISA; Hycult biotechnology, Netherlands).

### Histological analyses and acid phosphatase activity

Histologic analyses of lung lesions were performed at 3 days and 4 weeks after infection. Lungs embedded in paraffin for hematoxylin/eosin (HE) and Ziehl-Neelsen stainings. Lesions were evaluated on at least 3 lobe sections per animal. Lung sections were captured on a Zeiss Mirax Scan microscope system and analyzed using the Metamorph software as previously described [26]. For acid phosphatase staining, cryostat tissue sections from lung frozen in liquid nitrogen were used as previously described [27]. Signs of ROS production were evaluated by 8-hydroxy-2′-deoxyguanosine (8-OHdG) staining (1:50, JaICA, Shizuoka, Japan) as previously described [28].

### Ex-vivo recall responses of spleen cells and release of nitric oxide

Mice were infected with BCG, sacrificed at day 17 and spleen cells were prepared as previously described [29]. Cells were stimulated with either medium alone, living BCG (10^3^ CFU/well), or BCG culture protein extracts (17 μg/ml). After one, three and six days of treatment, medium was harvested for nitrite and TNF determination. Nitrite accumulation, as an indicator of NO production, was evaluated by Griess reagent (1% sulfanilamide and 0.1% naphtylethylenediamide in 2.5% phosphoric acid). TNF was determined in cell supernatants as described below.

### Evaluation of cytokines in lung homogenates

Lungs were collected at different time points after BCG injection and tissue homogenate was prepared [30]. Cytokines and chemokines were measured by ELISA (Ready&D System).

### Literature research

Literature research on CGD and mycobacterial infections was done from PubMed and Google Scholar with no limitations in time.

### Statistics

Parametric (*t*-tests) and non-parametric (One-way analysis and Kruskal–Wallis) tests were used. In the case of multiple comparisons, a two-way ANOVA test with Bonferroni correction was used.

---

**Supplementary Figure 1. Similar phenotypes in naïve mice.**

(A) Lung, liver and spleen histology (Hematoxylin and eosin staining) from wild-type, *Ncf1* mutant and *Ncf1* rescue mice without BCG infection. (B) Organ weight related to body weight of wild-type (n = 3), *Ncf1* mutant (n = 3) and *Ncf1* rescue (n = 3) mice without BCG infection. Magnifications were x100.

**Supplementary Figure 2. Lung damage in response to BCG infection in additional CGD mouse models.**

Lung histology (Hematoxylin and eosin staining) from NOX2 wild-type (left panels) and NOX2-deficient (right panels) mice, 3 days (A) and 4 weeks (B) after BCG infection. Three days post-infection, hemorrhagic pneumonia was observed in NOX-deficient mice. Four weeks after BCG infection, NOX2-deficient lungs show a massive inflammation, alveolar obstruction (B-b). Higher magnifications show massive infiltration of neutrophils only in NOX2-deficient lung (B-d). (C) *Ncf1* mutant mice with C57Bl/6 genetic background show also abscess of neutrophils, absent in respective wild-type mice. Magnifications were x100 and x1000.

**Supplementary Figure 3. Ex-vivo restimulation of splenocytes from BCG infected NOX2-deficient and wild-type mice.**

TNF and NO (nitric oxide) levels were evaluated in culture supernatant from splenocytes incubated with 10^3^ viable *M. bovis* BCG (A and C) or with antigens derived from *M. bovis* BCG (B and D). Values are shown as mean ± *SEM* (*n* = 4-5 mice per group, assayed in triplicate). * p < 0.05

**Supplementary Figure 4. Granuloma phenotype in additional CGD mouse models.**

Lung weight related to body weight of *Ncf1* mutant with C57Bl/6 genetic background and NOX2-deficient mice, as well as their respective controls 4 weeks post infection (**p < 0.01).

Representative hematoxylin and eosin (H&E) stained lung sections showing granulomas 4 weeks post infection. Magnifications were 200x.

